# The fluctuation-dissipation theorem and the discovery of distinctive off-equilibrium signatures of brain states

**DOI:** 10.1101/2024.04.04.588056

**Authors:** Juan Manuel Monti, Yonatan Sanz Perl, Enzo Tagliazucchi, Morten Kringelbach, Gustavo Deco

## Abstract

The brain is able to sustain many different states as shown by the daily natural transitions between wakefulness and sleep. Yet, the underlying complex dynamics of these brain states are essentially in non-equilibrium. Here, we develop a thermodynamical formalism based on the off-equilibrium extension of the fluctuation-dissipation theorem (FDT) together with a whole-brain model. This allows us to investigate the non-equilibrium dynamics of different brain states and more specifically to apply this formalism to wakefulness and deep sleep brain states. We show that the off-equilibrium thermodynamical signatures of brain states are significantly different in terms of the overall level of differential and integral violation of FDT. Furthermore, the framework allows for a detailed understanding of how different brain regions and networks are contributing to the off-equilibrium signatures in different brain states. Overall, this framework shows great promise for characterising and differentiating any brain state in health and disease.

## INTRODUCTION

A central problem in neuroscience is not only defining and characterising brain states but also discovering the underlying mechanisms. Brain states usually refer to distinct patterns of neural activity that occur during different conditions, such as wakefulness, sleep, and anesthesia [1–4]. Moving beyond such correlational definitions, Massimini and colleagues conducted a set of highly influential experiments using direct perturbations of the brain in different states to measure the dissipation of brain activity after perturbations [5–7]. This allowed them to assess the brain-wide spatiotemporal propagation of external stimulation using the Perturbational Complexity Index (PCI), which measures the amount of information contained in the amplitude of the average perturbation-evoked responses. PCI is computed as the Lempel-Ziv complexity in space and time of evoked EEG signals [5]. Thermodynamics has emerged as a promising framework for understanding non-equilibrium brain dynamics by describing the information flow by using the concepts of production entropy, the arrow of time, and irreversibility to measure the asymmetry resulting from the breaking of the detailed balance in different brain states [8–11].

Here, we show that the fluctuation-dissipation theorem (FDT) in non-equilibrium can lead to an even better understanding of the underlying mechanisms of the perturbational results from Massimini and colleagues. It has already been shown that the violation of the FDT can describe the different levels of non-equilibrium dynamics associated with different brain states through applying Onsager’s regression principle and observing the evolution of the different brain areas following a perturbation [12]. More generally, here we adopt the off-equilibrium extension of the FDT [13–15], to develop a model-based analytical framework in which we only use data from the unperturbed system to capture deviations from equilibrium dynamics arising in different brain states. These deviations directly serve as distinctive markers of brain states.

We used this new off-equilibrium FDT framework on empirical human neuroimaging data of wakefulness and deep sleep. In contrast to Massimini’s perturbational framework, we do not need to empirically perturb the brain but can directly show significantly different off-equilibrium thermodynamical signatures of brain states in terms of the overall level of differential and integral violation of FDT. Furthermore, the framework also allows for a more detailed understanding of how different brain regions and networks contribute to the off-equilibrium signatures in different brain states.

## OFF-EQUILIBRIUM FDT

A way of expressing the FDT is through the relation between the linear response function and the auto-correlation function. The linear response function of an observable *x* of the system to a small enough perturbation *h* is given by

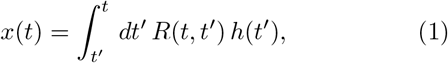

whereas the auto-correlation of *x* is defined as

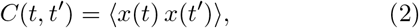

here ⟨…⟩ states for the ensemble average. The FDT states that, if a system is in equilibrium, the linear response function is related to the auto-correlation function of the unperturbed observable by

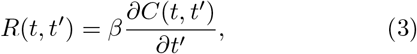

where *t*^′^ ≤ *t*, and *β* = 1*/T*, being *T* the temperature of the thermal bath. However, if the system is out of equilibrium, the generalized relation between linear response and auto-correlation is given by (see [13])

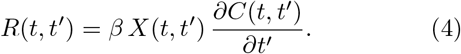

In (4) *X*(*t, t*^′^) is called the Fluctuation-Dissipation Ratio and characterizes the approach to equilibrium and measures the deviation from FDT. Also, as introduced in [14], a way to measure the separation from FDT, and therefore from equilibrium, is through the Differential Violation *V* (*t, t*^′^), and Integral Violation *I*(*t, t*^′^) of FDT given by

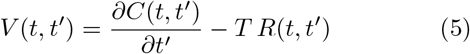

and

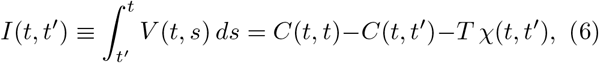

respectively, where 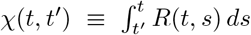 is the integrated response function, or dynamic susceptibility. For systems in equilibrium *X* = 1, *V* = 0, and *I* = 0.

If we now consider a system evolving with the Langevin equation

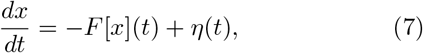

where *F* is a deterministic function and *η*(*t*) is a zero-mean Gaussian noise with auto-correlation ⟨*η*(*t*)*η*(*t*^′^)⟩ = 2*T δ*(*t*−*t*^′^), we can also obtain the linear response through the expressions (see [13])

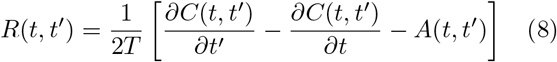

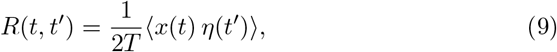

where *A*(*t, t*^′^) is the so called asymmetry

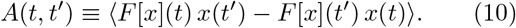

Therefore, if a system is governed by Langevin dynamics, it is possible to determine *C*(*t, t*^′^) and then *R*(*t, t*^′^) with equations (8) or (9), and evaluate its distance from FDT using *X*(*t, t*^′^), *V* (*t, t*^′^) and *I*(*t, t*^′^). The degree of non-equilibrium determined by the distance from FDT could then be a marker of brain states (e.g. consciousness, sleep, coma). To prove this hypothesis, we use empirical human neuroimaging data (fMRI, functional Magnetic Resonance Imaging) for wakefulness and deep sleep, in order to fit a whole-brain model that can be written as a Langevin equation to simulate data from which the linear response *R*(*t, t*^′^) can be calculated using equations (8) or (9) to further determine *X*(*t, t*^′^), *V* (*t, t*^′^) and *I*(*t, t*^′^).

## WHOLE-BRAIN MODEL

Previous research has proposed that different brain states are characterised by different dynamical regimes [16, 17]. As a matter of fact, it was shown that at the macroscopic and microscopic scales, unconscious brain states are dominated by synchronous activity [16–20]. In contrast, conscious states are characterised by asynchronous dynamics [17, 21, 22]. Aligned with this, we consider a whole-brain model consisting of *N* coupled brain regions (nodes) where the local dynamic of each brain region is described using a Hopf bifurcation in its normal form (also known as a Stuart-Landau oscillator) (see [23, 24]). With this, each node can present an asynchronous noisy state or display coherent oscillations when being below or above a given threshold, respectively. Then, each node *j* of the brain network is modeled by a normal Hopf bifurcation as:

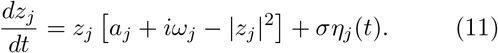

In (11) *a*_*j*_ is the so-called bifurcation parameter, *ω*_*j*_ the node frequency, and *ση*_*j*_(*t*) is a zero-mean additive Gaussian noise with standard deviation *σ*. We then consider the coupling of the network nodes through the anatomical network as given by the general effective connectivity matrix (GEC) 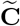 (see SI Appendix). Hence, the whole-brain dynamics is modeled via the following set of coupled equations:

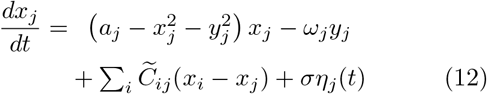

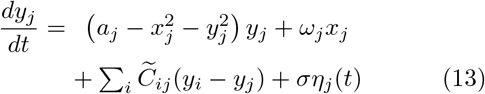

where *x*_*j*_ is the simulated signal for each node *j*. Then, the whole-brain model given by equations (12) and (13) can be written as a set of coupled Langevin equations.

### Model parameters. Fitting of the Hopf whole-brain model to empirical data

The empirical fMRI BOLD data, to which the whole-brain model is fitted (see Fig. 1), is constituted of two cohorts corresponding to wakefulness and deep sleep (N3 stage of sleep) brain states, each composed of 15 individual scans (subjects), with *N* = 90 nodes and *n*_*F*_ = 180 time frames. The only fixed parameter in the model is the local bifurcation parameter which is set to the value *a*_*j*_ = −0.02 for all subjects and the different brain states considered. As stated in [25], this corresponds to all the oscillators being at the brink of the bifurcation. The node frequencies *ω*_*j*_, for each brain state, are determined as the peak of the mean power spectrum of the empirical time series [24]. Finally, the standard deviation *σ* of the Gaussian noise is obtained by searching for the value of *σ* that gives the least difference between the empirical and simulated matrices defined as

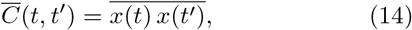

here the bar means performing the average over all the subjects in each given brain state (see SI Appendix).

**FIG. 1.**
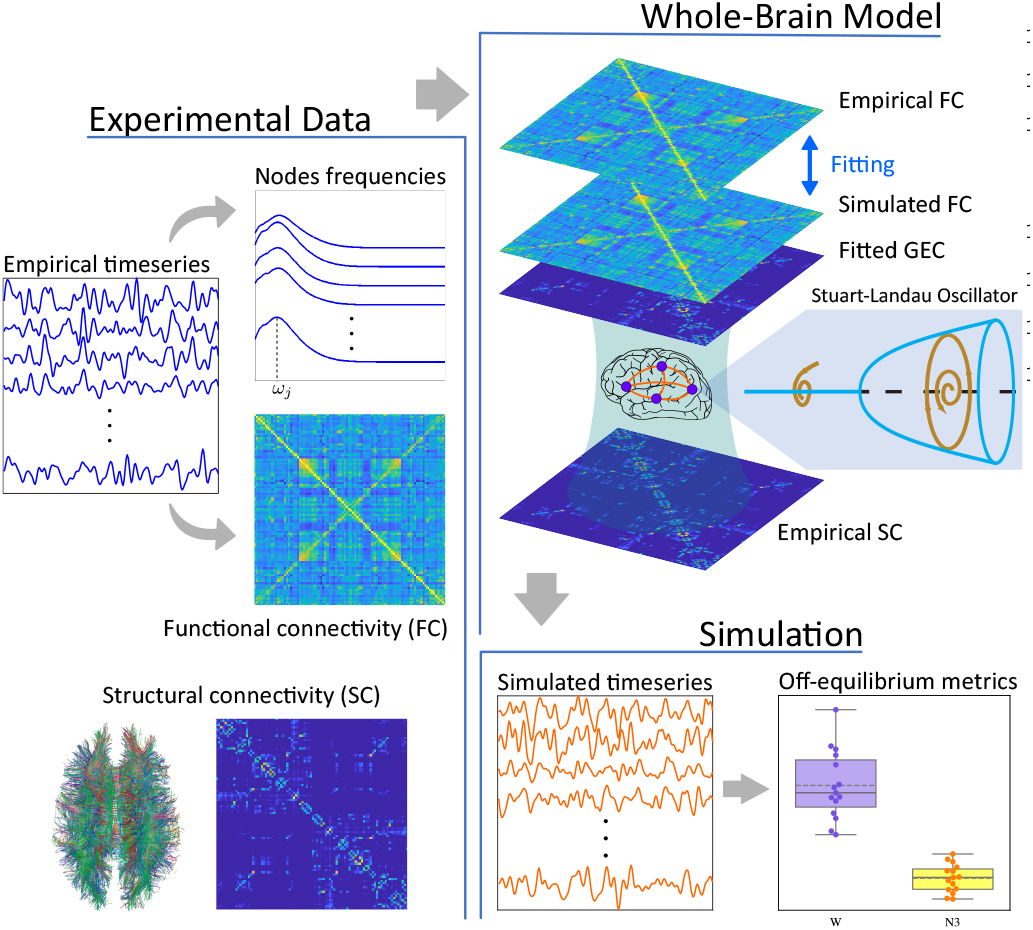
Model-based non-equilibrium FDT. Using empirical data we link anatomical connectivity and functional brain connectivity in order to fit a Hopf whole-brain model to generate simulated data that satisfies Eq. (7). The simulated data is then used to determine the distance from equilibrium using Eqs. (15), (16), and (17).

## APPLYING THE FORMALISM TO WAKEFULNESS AND DEEP SLEEP BRAIN STATES

Once the whole-brain model is fitted to each subject, simulated time series can be obtained, from which auto-correlation (Eq. (2), and the linear response (equations (8) or (9) can be calculated. For each subject, and each node, we can obtain three *n*_*F*_ × *n*_*f*_ sub-diagonal matrices (X, V, and I) and define three metrics to determine the distance from equilibrium (see Fig. 2), *i*.*e*. the deviation from FDT, namely:

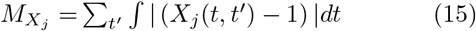

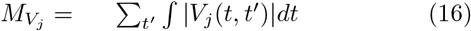

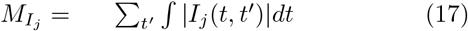

each of these metrics can give a measure of the distance from the equilibrium of the brain region *j*. Then, for each subject, we can take the average over all nodes as a more global indicator of distance from equilibrium.

**FIG. 2.**
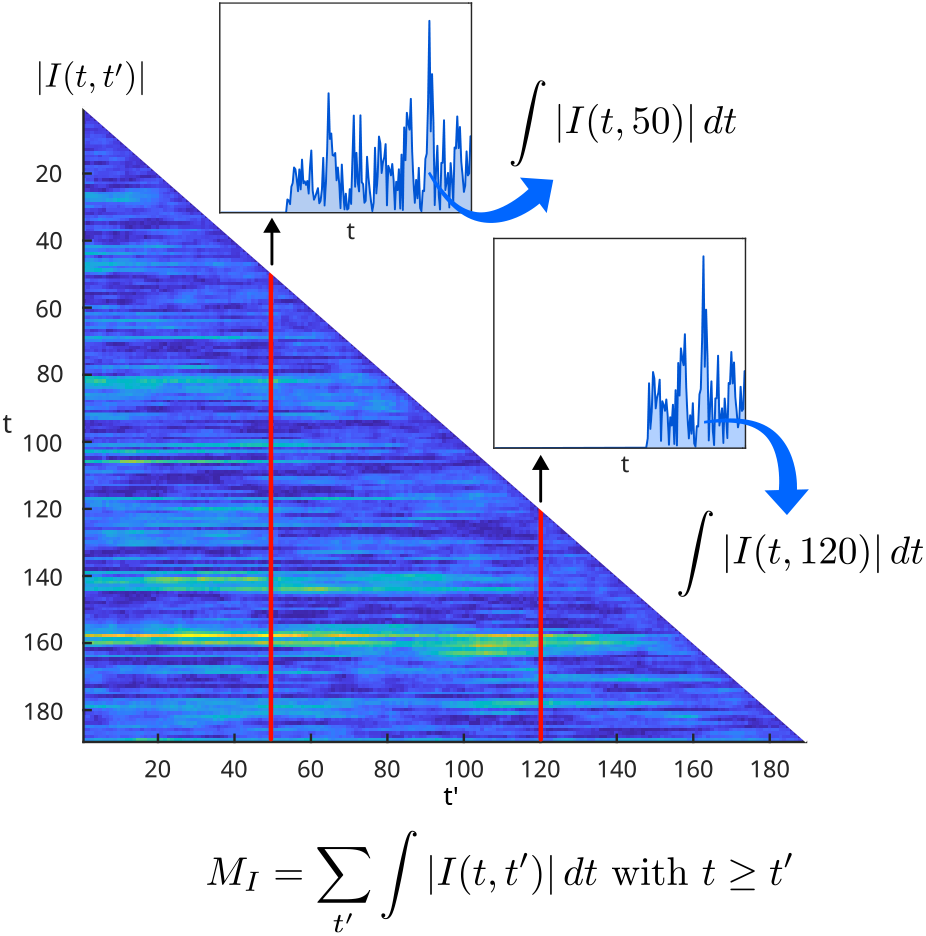
Calculations of metrics. Typical sub-diagonal matrix obtained for |*I*(*t, t*^′^)| for one brain region. For each time-step *t*^′^ (showing *t*^′^ = 50 and *t*^′^ = 120 as examples), we calculate the mean over *t* (with *t ≥ t*^′^) and the sum over all *t*^′^ to obtain the different metrics.

## BRAIN STATES CHARACTERIZATION

In Fig. 3, we show the average over all the brain regions for each subject in the wakefulness and deep sleep groups. The ensemble average was determined by performing 10.000 simulations for each subject and each brain state. A discussion over the convergence of the metrics with respect to the number of simulations can be found in the SI Appendix. In general, all three metrics show a higher departure from equilibrium for the wakefulness group, with statistical significance *p <* 1*e*^−3^ for *M*_*X*_, and *p <* 1*e*^−5^ for *M*_*V*_ and *M*_*I*_ (calculated with a Wilcoxon rank-sum test) between the distributions. Also for the Differential and Integral Violations of FDT, there is no overlapping between the swarmplots which shows high distinguishability between states.

**FIG. 3.**
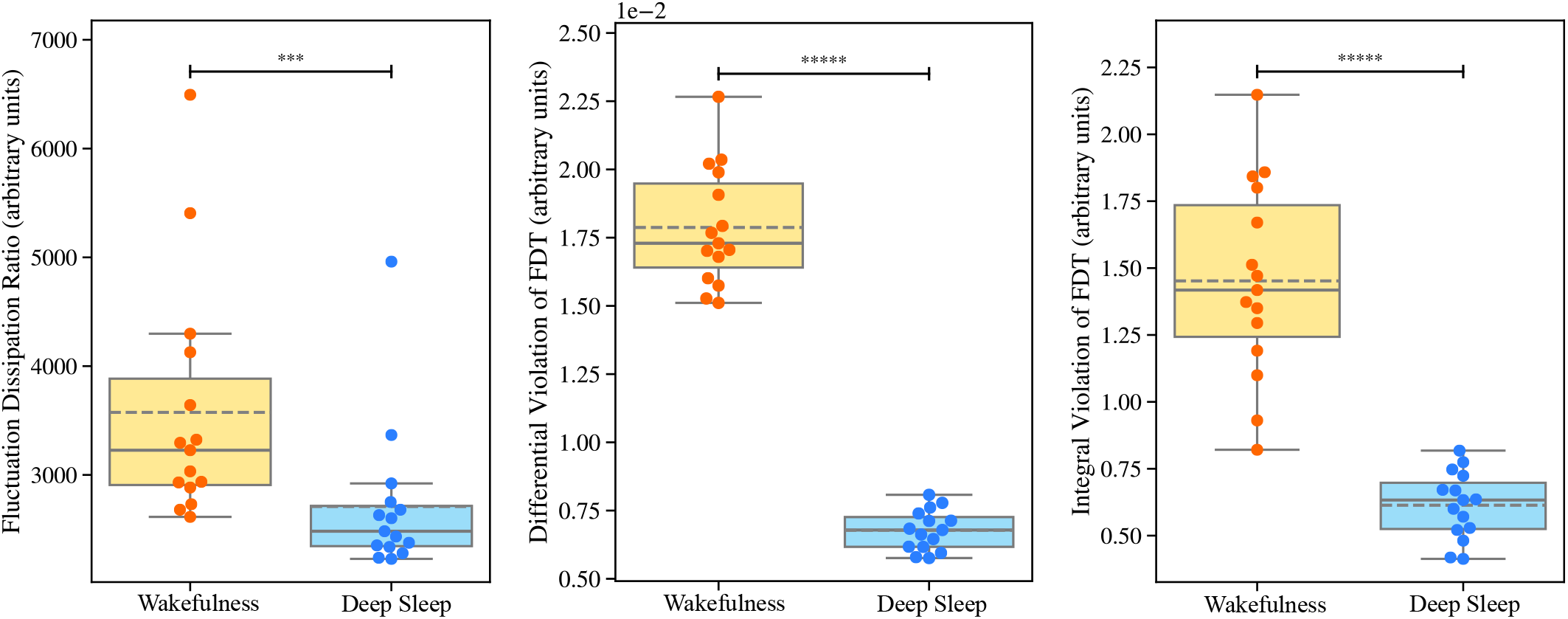
Non-equilibrium characterization of brain states: metrics across subjects. Deviation from FDT, i.e. distance from equilibrium calculated as the average for all nodes for each subject using the metrics *M*_*X*_ (left), *M*_*V*_ (center), and *M*_*I*_ (right). These metrics are defined by the Fluctuation Dissipation Ratio (Eq. (15)), the Differential Violation of FDT (Eq. (16)), and the Integral Violation of FDT (Eq. (17)), respectively. Solid lines represent the median of the distributions, whilst the dashed lines indicate their mean.

The distribution plots in all cases show more spreading in the wakefulness group, whereas the deep sleep distribution appears more clustered. This aspect may reflect the fact that the deep-sleep state has less inter-subject variability than wakefulness.

Of the three given metrics the one that gives more reliable results is the *M*_*I*_ as it does not depend on the numerical derivation of a noisy function. Instead, it involves an integration, which is numerically much more reliable. Also, it shows the fastest convergence with respect to the number of simulations (see the SI Appendix).

On the other hand, if instead of performing an average over nodes, a mean over subjects is considered, we can obtain the deviation from equilibrium for each brain region, which is shown in Fig. 4. This could be used to determine the intrinsic hierarchy of brain regions that characterize different brain states.

**FIG. 4.**
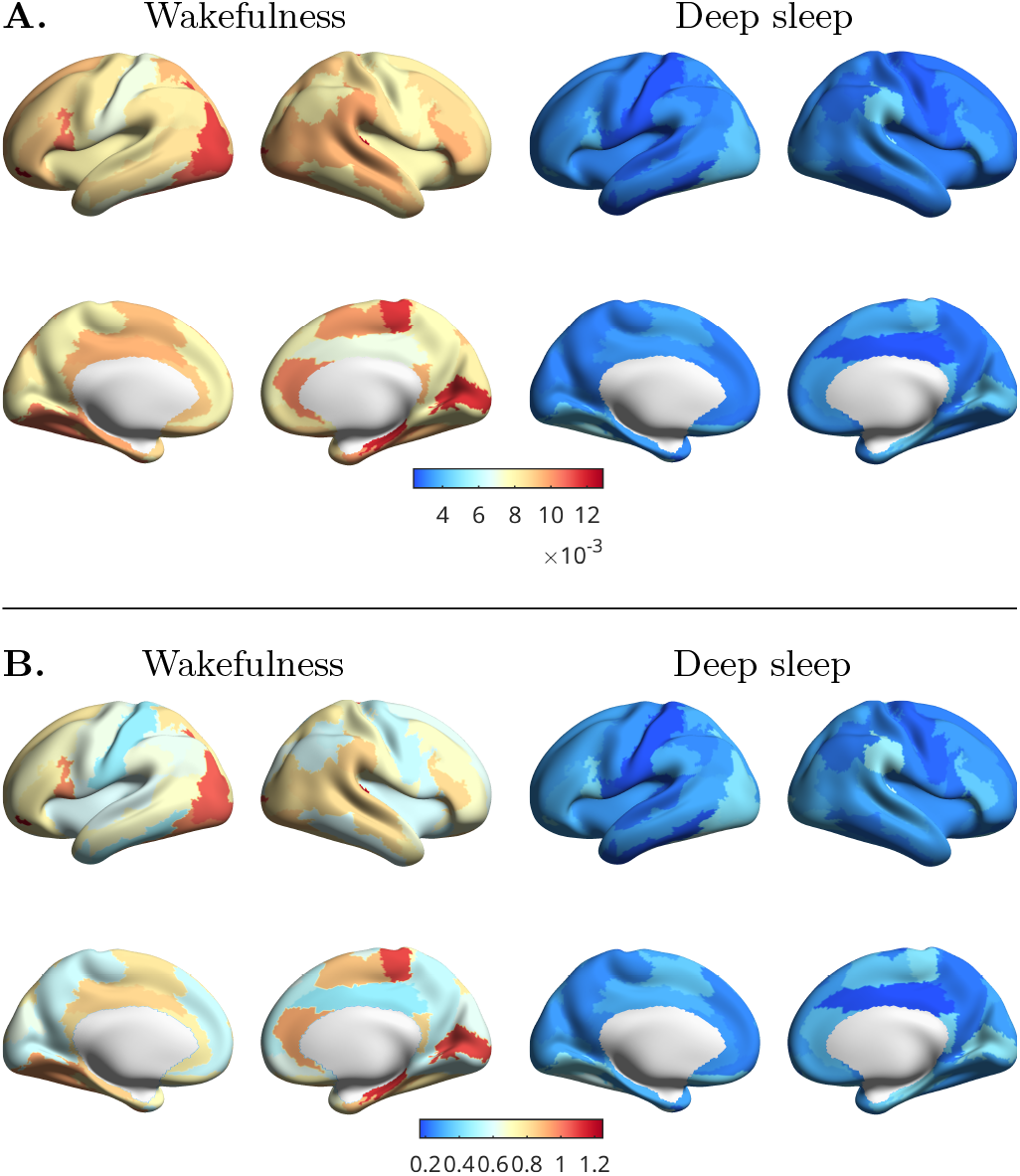
Non-equilibrium characterization of brain states: metrics across brain areas. Here we also consider averaging over subjects instead of nodes, hence the level of non-equilibrium can be studied for each region on the wakefulness (left plots) and deep-sleep states (right plots). **A**. Differential Violation of FDT (Eq. (16)). **B**. Integral Violation of FDT (Eq. (17)).

## DISCUSSION

We have presented a model-based formalism using the Hopf whole-brain model together with the off-equilibrium generalization of the FDT theorem presented in [13], with which the unperturbed simulated timeseries can be used to determine the departure from equilibrium of different brain states. The results show that there is a clear distinction between the wakefulness and deep sleep states based on their distance from equilibrium determined by the metrics derived from the Fluctuation-Dissipation Ratio [13], the Differential and the Integral Violations of FDT [14]. More precisely, we found that the deviations from equilibrium were larger in the wakefulness state than in deep sleep. These findings are in agreement with those from [9] where the level of non-reversibility of different brain states and cognitive tasks are evaluated in electrocorticography data from non-human primates showing larger levels of irreversibility for wakefulness than for deep sleep.

As stated above, there is a correspondence between the temperature of the thermal bath in which the system evolves [13] and the standard deviation of the noise in the Hopf model. This would indicate that, at least in the model, each brain state also has a characteristic noise. Following the analogy, we may think that each brain state is a different thermal bath in which the system (brain) evolves. This also sheds light on the role of noise in the model, not only present to generate transitions between the noisy and oscillatory regimes of the Hopf bifurcation but also playing a more fundamental role in defining the brain state.

The off-equilibrium FDT formalism presented here has the advantage that only data from the unperturbed simulated system is needed to determine the metrics that define the distance from equilibrium. This avoids the need of perturbing the system, whether experimentally or in-silico.

Summing up, here we provide a model-based framework that can be used to evaluate the level of non-equilibrium by determining the degree of violation of FDT through the metrics provided. This allows us to characterize distinct brain states, namely wakefulness and deep sleep. All the metrics show a larger level of non-equilibrium in the wakefulness state than in deep sleep. This is in clear alignment with the idea that non-equilibrium brain dynamics is a general feature of conscious states [11].

Future research could extend this investigation by considering other brain states such as different degrees of comatose states, e.g. minimally conscious state and unresponsiveness wakefulness syndrome, diverse cognitive tasks, and different stages of neurodegenerative diseases and stroke.

## ACKNOWLEDGMENTS

J.M.M. acknowledges financial support from CONICET. G.D. was supported by several sources: NEMESIS project (ref. 101071900) funded by the EU ERC Synergy Horizon Europe; AGAUR research support grant (ref. 2021 SGR 00917) funded by the Department of Research and Universities of the Generalitat of Catalunya; project PID2022-136216NB-I00 financed by the MCIN /AEI /10.13039/501100011033 / FEDER, UE., the Ministry of Science and Innovation, the State Research Agency and the European Regional Development Fund. Y.S.P is supported by European Union’s Horizon 2020 research and innovation program under the Marie Sklodowska-Curie grant 896354.

