## Supplementary Information for "The fluctuation-dissipation theorem and the discovery of distinctive off-equilibrium signatures of brain states"

#### 1. PARCELLATION

Both datasets (for wakefulness and deep-sleep) used timeseries from the AAL90 parcellation with a total of 90 cortical regions (see [1]).

#### 2. STRUCTURAL CONNECTIVITY

The structural connectivity data was obtained from the March 2017 public data release from the Human Connectome Project (HCP).

##### 2.a. 3T structural data

The HCP structural data were acquired using a customized 3 Tesla Siemens Connectom Skyra scanner with a standard Siemens 32-channel RF-receive head coil. For each participant, at least one 3D T1w MPRAGE image and one 3D T2w PACE image were collected at 0.7 mm isotropic resolution.

##### 2.b. 3T diffusion MRI

In order to reconstruct a high-quality structural connectivity (SC) matrix for constructing the whole-brain model (using the AAL90 parcellation), we obtained multi-shell diffusion-weighted imaging data from 32 participants from the HCP database (scanned for approximately 89 minutes). The acquisition parameters are described in detail on the HCP website [2]. We estimated the connectivity using the method described by Horn and colleagues [3]. Briefly, the data was processed using a generalized q-sampling imaging algorithm implemented in DSI studio (http://dsi-studio.labsolver.org). Segmentation of the T2-weighted anatomical images produced a white-matter mask and co-registering the images to the b0 image of the diffusion data using SPM12. In each HCP participant, 200,000 fibres were sampled within the white-matter mask. Fibres were transformed into MNI space using Lead-DBS [4]. The methods used the algorithms for false-positive fibres shown to be optimal in recent open challenges [5, 6]. The risk of false positive tractography was reduced in several ways. Most importantly, this used the tracking method achieving the highest (92%) valid connection score among 96 methods submitted from 20 different research groups in a recent open competition [5].

### 3. HUMAN SLEEP DATA: ACQUISITION AND PRE-PROCESSING

#### 3.a. Ethics

Written informed consent was obtained, and the study was approved by the ethics committee of the Faculty of Medicine at the Goethe University of Frankfurt, Germany.

#### 3.b. Participants

We used fMRI- and PSG data from 15 participants taken from a larger database that reached all four stages of PSG [7]. Exclusion criteria focused on the quality of the concomitant acquisition of EEG, EMG, fMRI, and physiological recordings.

#### 3.c. Acquisition and pre-processing of fMRI and polysomnography data

Neuroimaging fMRI was acquired on a 3 T system (Siemens Trio, Erlangen, Germany) with the following settings: 1505 volumes of T2*-weighted echo planar images with a repetition time (TR) of 2.08 s, and an echo time of 30 ms; matrix $64\times64$, voxel size $3\times3\times2$ mm$^3$, distance factor 50%, FOV 192 mm$^2$.

The EPI data were realigned, normalised to MNI space, and spatially smoothed using a Gaussian kernel of 8 mm$^3$ FWHM in SPM8 (http://www.fil.ion.ucl.ac.uk/spm/). Spatial downsampling was then performed to a $4\times4\times4$ mm resolution. From the simultaneously recorded ECG and respiration, cardiac- and respiratory-induced noise components were estimated using the RETROICOR method [8], and together with motion parameters these were regressed out of the signals. The data were temporally band-pass filtered in the range 0.008-0.08 Hz using a sixth-order Butterworth filter. We extracted the timeseries in the AAL90 parcellation [1].

Simultaneous PSG was performed through the recording of EEG, EMG, ECG, EOG, pulse oximetry, and respiration. EEG was recorded using a cap (modified BrainCapMR, Easycap, Herrsching, Germany) with 30 channels, of which the FCz electrode was used as reference. The sampling rate of the EEG was 5 kHz, and a low-pass filter was applied at 250 Hz. MRI and pulse artifact correction were applied based on the average artifact subtraction method [9] in Vision Analyzer2 (Brain Products, Germany). EMG was collected with chin and tibial derivations, and as the ECG and EOG recorded bipolarly at a sampling rate of 5 kHz with a low-pass filter at 1 kHz. Pulse oximetry was collected using the Trio scanner, and respiration with MR-compatible devices (BrainAmp MR+, BrainAmp ExG; Brain Products, Gilching, Germany).

Participants were instructed to lie still in the scanner with their eyes closed and relax. Sleep classification was performed by a sleep expert based on the EEG recordings in accordance with the AASM criteria (2007). Results using the same data and the same pre-processing have previously been reported [7].

#### 4. BRAIN NETWORK MODEL

The brain network model consists of 90 coupled brain areas (nodes) derived from the AAL90 parcellation. The global dynamics of the brain network model used here result from the mutual interactions of local node dynamics coupled through the underlying general effective connectivity $\widetilde{C}_{ij}$. The local dynamics of each individual node are described by the normal form of a supercritical Hopf bifurcation (see Figure S1), which is able to describe the transition from asynchronous noisy behavior to full oscillations. Thus, in complex coordinates, each node $j$ is described by the following equation:

$$\frac{dz_j}{dt} = z\left[a_j + i\omega_j - |z_j|^2\right] + \sigma\eta_j(t) \tag{S1}$$

where

$$z_j = \rho_j \exp\left(i\,\theta_j\right) = x_j + i\,y_j \tag{S2}$$

and $\eta_j(t)$ is additive Gaussian noise with standard deviation $\sigma$. This normal form has a supercritical bifurcation at $a_j = 0$ so that for $a_j < 0$ the local dynamics have a stable fixed point at $z_j = 0$ (which due to the additive noise corresponds to a low activity asynchronous state) and for $a_j > 0$ there exists a stable limit cycle oscillation with frequency $f_j = \omega_j/2\pi$. We insert Equation (S2) in Equation (S1) and separate the real part in Equation (S3) and the imaginary part in Equation (S4):

$$\frac{dx_j}{dt} = \left[a_j - x_j^2 - y_j^2\right] x_j - \omega_j y_j + \sigma\eta_j(t) \tag{S3}$$

$$\frac{dy_j}{dt} = \left[a_j - x_j^2 - y_j^2\right] y_j + \omega_j x_j + \sigma\eta_j(t) \tag{S4}$$

We then consider the coupling of the network nodes through the anatomical network as given by the general effective connectivity matrix $\widetilde{\mathbf{C}}$. Thus, the whole-brain dynamics is defined by the following set of coupled equations:

$$\frac{dx_j}{dt} = \left(a_j - x_j^2 - y_j^2\right) x_j - \omega_j y_j + \textstyle\sum_i \widetilde{C}_{ij}(x_i - x_j) + \sigma\eta_j(t) \tag{S5}$$

$$\frac{dy_j}{dt} = \left(a_j - x_j^2 - y_j^2\right) y_j + \omega_j x_j + \textstyle\sum_i \widetilde{C}_{ij}(y_i - y_j) + \sigma\eta_j(t) \tag{S6}$$

We couple the equations using the common difference coupling, which approximates the simplest (linear) part of a general coupling function. These equations are valid in the weakly coupled oscillator limit, in which the coupling preserves the periodic orbit of the uncoupled oscillators. We then model with the variables $x_j$ the BOLD signal of each node $j$.

##### 4.a. Nodes frequencies

The intrinsic frequencies $\omega_j$ (which lie in the 0.008–0.08 Hz band) were estimated from the data as the averaged peak frequencies of the narrowband blood-oxygen-level-dependent

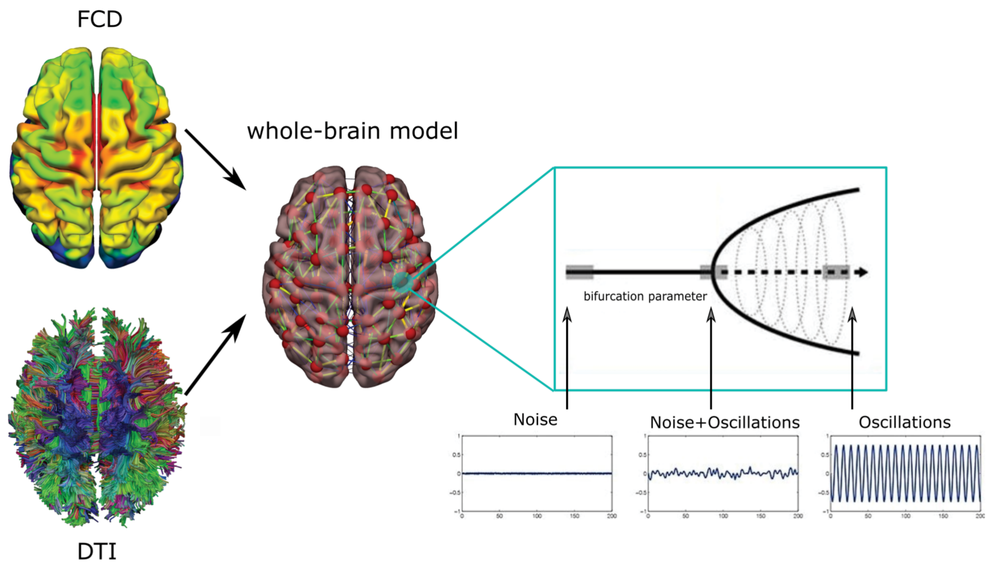


**Fig. S1. Hopf whole-brain model.** Each brain region is modeled as a supercritical Hopf bifurcation. The empirical functional and structural connectivities are used in order to obtain the coupling between the nodes.

(BOLD) signals of the different brain regions. For $a_j > 0$, the local dynamics settle into a stable limit cycle, producing self-sustained oscillations with frequency $\omega_j/(2\pi)$. For $a_j < 0$, the local dynamics present a stable spiral point, producing damped or noisy oscillations in the absence or presence of noise, respectively.

##### 4.b. General Effective Connectivity

For the optimization of the coupling matrix $\widetilde{\mathbf{C}}$, we estimate it numerically by using a pseudo-gradient descent procedure [10, 11]. Specifically, we fit $\widetilde{\mathbf{C}}$ such that the model optimally reproduces the empirically measured covariances $\mathbf{FC}^{\text{empirical}}$ (i.e., the normalized covariance matrix of the functional neuroimaging data) and the empirical time-shifted covariances $\mathbf{FS}^{\text{empirical}}(\tau)$, where $\tau$ is the time lag. We selected the parameter $\tau$, which led to a decrease in the averaged autocorrelation. These normalized time-shifted covariance matrices are generated by taking the shifted covariance matrix $\mathbf{KS}^{\text{empirical}}(\tau)$ and dividing each pair $(i, j)$ by $\sqrt{KS_{ii}^{\text{empirical}}(0)KS_{jj}^{\text{empirical}}(0)}$. Note that these normalized time-shifted covariances break the symmetry of the couplings and thus improve the level of fitting [12]. Using a heuristic pseudo-gradient algorithm, we proceeded to update the $\widetilde{\mathbf{C}}$ until the fit was fully optimized. More specifically, the updating uses the following form:

$$\widetilde{C}_{ij} = \widetilde{C}_{ij} + \alpha \left( FC_{ij}^{\text{empirical}} - FC_{ij}^{\text{model}} \right) + \delta \left( FS_{ij}^{\text{empirical}}(\tau) - FS_{ij}^{\text{model}}(\tau) \right) \tag{S7}$$

where $FS_{ij}^{\text{model}}(\tau)$ is defined similar to $FS_{ij}^{\text{empirical}}(\tau)$. Details on the computation of the **KS** matrix can be found in [13]. The model was run repeatedly with the updated $\widetilde{\mathbf{C}}$ until the fit converged towards a stable value. We initialized $\widetilde{\mathbf{C}}$ using the anatomical connectivity **C**. The **C** matrix denotes the density of fibers between cortical area $i$ and $j$ (obtained with probabilistic tractography from dMRI) and only updated known existing connections from this matrix (in either hemisphere). We used $\alpha = \delta = 1e^{-5}$ and continued until the algorithm converged. For each iteration, we compute the model results as the average over as many simulations as there are subjects. Overall, we use the term Generative Effective Connectivity (GEC) for the optimized $\widetilde{\mathbf{C}}$ [14].

##### 4.c. Standard deviation of the noise

The standard deviation of the noise was determined by varying the value of $\sigma$ and calculating the difference between the empiric and simulated $\overline{\mathbf{C}}$ matrices defined as:

$$\overline{C_j}(t,t') = \frac{1}{N_{\text{sub}}} \sum_{n=1}^{N_{\text{sub}}} x_{n,j}(t) x_{n,j}(t') \tag{S8}$$

where $N_{\text{sub}}$ is the number of subjects, $j$ denotes the node, and $x(t)$ is the previously normalized empiric or simulated time-serie. The normalization was performed by dividing each time-serie by its standard deviation (z-score normalization).

In the case of the simulated $\overline{\mathbf{C}}$ matrices, the $N_{\text{sub}}$ simulations to calculate (S8) were performed using the group-average GEC matrix, which is the GEC matrix optimized to fit the mean effective connectivity. For each brain region, the difference between matrices was calculated as

$$d_j = \frac{1}{n_f^2} \sum_t \sum_{t'} \left[ \overline{C_j^{\text{emp}}}(t,t') - \overline{C_j^{\text{sim}}}(t,t') \right]^2 \tag{S9}$$

where $n_f$ is the number of time frames. Then, a single value to characterize the distance between matrices is obtained by averaging over all nodes:

$$D = \frac{1}{N} \sum_{j=1}^{N} d_j \tag{S10}$$

Finally, for each $\sigma$ value this process was repeated 100 times in order to increase statistics.

The optimal values for $\sigma$, for each brain state, are shown in Figure S2, where it can be seen that for the W (wakefulness) state we find $\sigma_W = 0.12$ and for N3 (deep sleep) $\sigma_{N3} = 0.06$.

#### 5. DIFFERENT WAYS OF CALCULATING $R(t,t')$

In [15] it is stated that if a system verifies a Langevin dynamic,

$$\frac{dx}{dt} = -F[x](t) + \eta(t), \tag{S11}$$

where $F$ is a deterministic function and $\eta(t)$ is a zero-mean Gaussian noise with autocorrelation $\langle \eta(t)\eta(t') \rangle = 2T\,\delta(t-t')$ then the linear response function can be determined

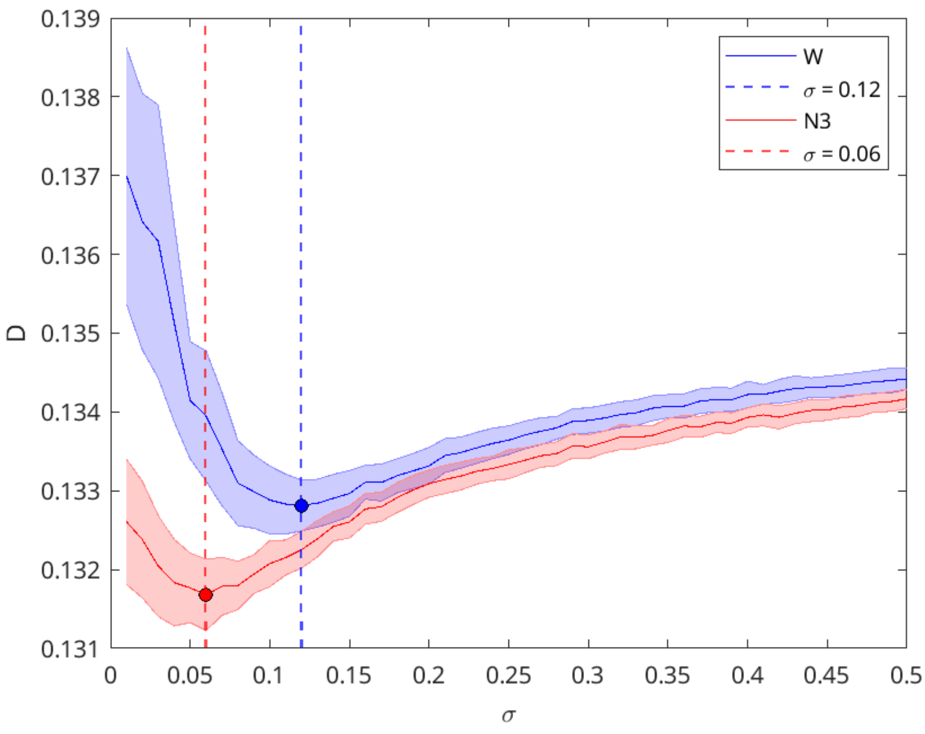


**Fig. S2. Optimal noise standard deviation.** Difference of the empiric and simulated $\overline{C}$ matrices as a function of the standard deviation $\sigma$.

as:

$$R(t,t') = \frac{1}{2T}\left[\frac{\partial C(t,t')}{\partial t'} - \frac{\partial C(t,t')}{\partial t} - A(t,t')\right] \quad \text{(S12)}$$

$$R(t,t') = \frac{1}{2T}\langle x(t)\,\eta(t')\rangle \quad \text{(S13)}$$

where $A(t,t')$ is the so called asymmetry

$$A(t,t') \equiv \langle F[x](t)\,x(t') - F[x](t')\,x(t)\rangle. \quad \text{(S14)}$$

Expressions (S12) and (S13) are equivalent but in practice, calculating the numerical derivative of a noisy function is not at all a trivial task. All the results presented in the main text are calculated considering the expression (S13) for the linear response function. However, we also show the results obtained with the expression (S12). The numerical derivatives were calculated using the `derivative` code https://github.com/tamaskis/derivative-MATLAB which calculates the derivative at lower (upper) bound using forward (backward) difference and approximates derivatives at all other nodes using central differences.

Figures S3 show results similar to the ones presented in the main text, but the Fluctuation Dissipation Ratio shows a large spreading and less statistical significance, due to the numerical calculation of the derivative of $C(t,t')$.

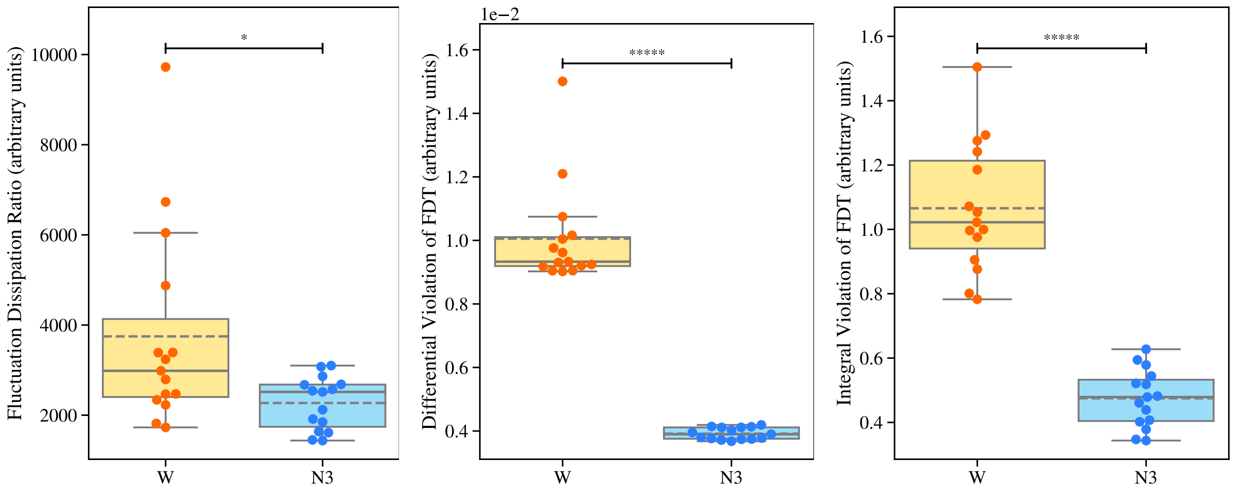


**Fig. S3. Non-equilibrium characterization of brain states: metrics across subjects**. Deviation from FDT, i.e. distance from equilibrium calculated as the average for all nodes using the metric $M_X$ related to the Fluctuation Dissipation Ratio (**A**), $M_V$ related with the Differential Violation of FDT (**B**), and $M_I$ related with the Integral Violation of FDT (**C**), where the linear response is calculated with expression (S12).

#### 6. CONVERGENCE OF THE METRICS

Determining the metrics presented in the main text involves calculating an ensemble average, which consists of the mean over a *large enough* number of system evolutions. In order to set this number with appropriate criteria, we study the convergence of each metric with respect to the number of simulations. It is clear that the $M_X$ metric has the worst convergence showing a large discrepancy between the distribution mean and median due to the quotient of the transition amplitude with the numerical calculation of the autocorrelation derivative. As for the $M_V$ and $M_I$ metrics, the difference between the mean and the median is almost negligible, and they show a steadily decreasing tendency with the increasing number of simulations, being the latter the one presenting a clear convergence for a number of simulations larger than 1000, whereas $M_V$ seems to converge for a number of simulations of around 10000. This might be due to the fact that the $M_I$ metric does not involve any numerical differentiation. Instead, in its calculation, a numerical integration is required, which makes it a far more stable calculation. These findings make $M_I$ the more reliable metric of the three presented in this work.

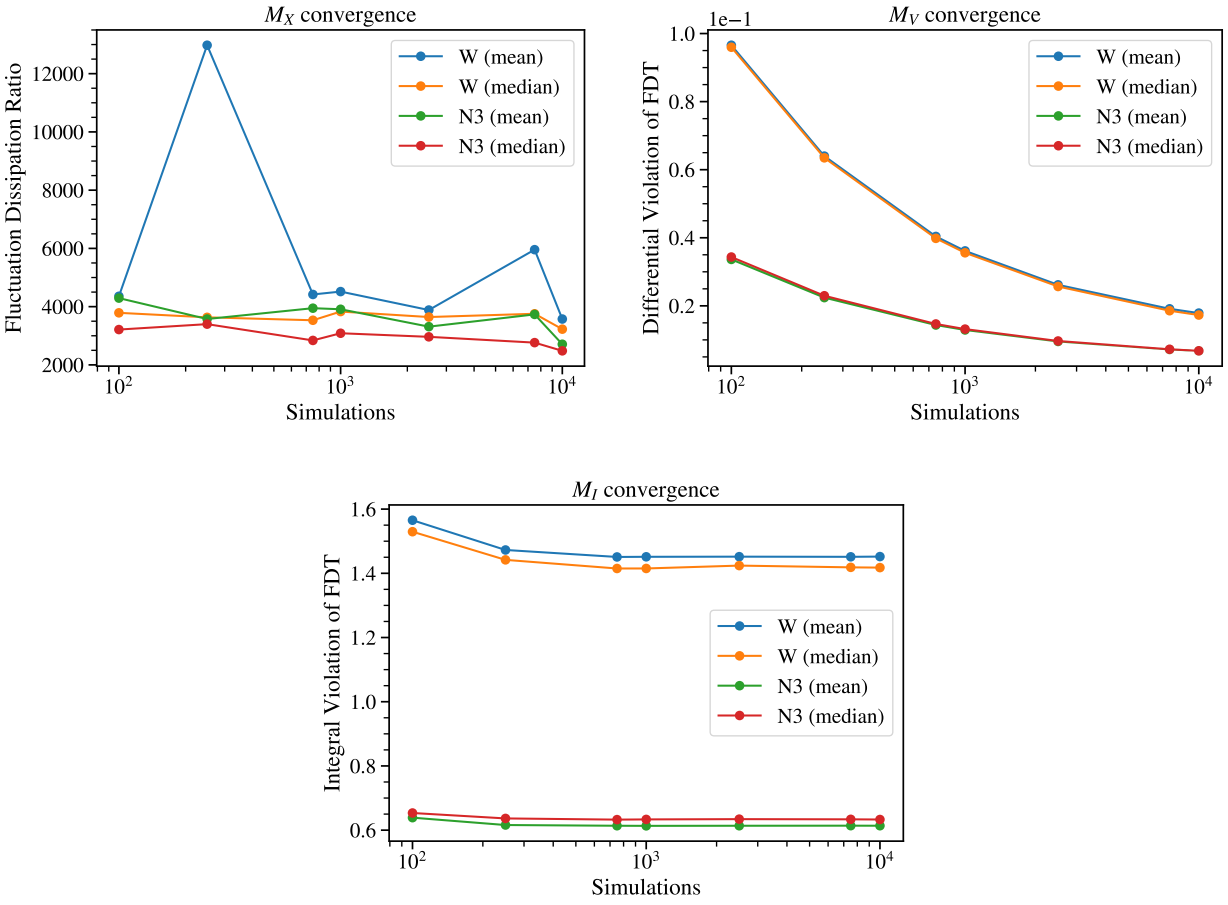


**Fig. S4. Convergence of the metrics.** We plotted the mean over all the subjects for both brain states for different numbers of simulations.

3. A. Horn, W.-J. Neumann, K. Degen, *et al.*, “Toward an electrophysiological “sweet spot” for deep brain stimulation in the subthalamic nucleus,” Hum. Brain Mapp. **38**, 3377–3390 (2017).
4. A. Horn and F. Blankenburg, “Toward a standardized structural–functional group connectome in mni space,” NeuroImage **124**, 310–322 (2016).
5. K. H. Maier-Hein, P. F. Neher, J.-C. Houde, *et al.*, “The challenge of mapping the human connectome based on diffusion tractography,” Nat. Commun. **8**, 1349 (2017).
6. K. G. Schilling, A. Daducci, K. Maier-Hein, *et al.*, “Challenges in diffusion mri tractography – lessons learned from international benchmark competitions,” Magn. Reson. Imaging **57**, 194–209 (2019).
7. A. B. A. Stevner, D. Vidaurre, J. Cabral, *et al.*, “Discovery of key whole-brain transitions and dynamics during human wakefulness and non-rem sleep,” Nat. Commun. **10**, 1035 (2019).
8. G. H. Glover, T.-Q. Li, and D. Ress, “Image-based method for retrospective correction of physiological motion effects in fmri: Retroicor,” Magn. Reson. Medicine **44**, 162–167 (2000).
9. P. J. Allen, G. Polizzi, K. Krakow, *et al.*, “Identification of eeg events in the mr scanner:

The problem of pulse artifact and a method for its subtraction," NeuroImage **8**, 229–239 (1998).

10. M. Gilson, R. Moreno-Bote, A. Ponce-Alvarez, *et al.*, "Estimation of directed effective connectivity from fmri functional connectivity hints at asymmetries of cortical connectome," PLOS Comput. Biol. **12**, 1–30 (2016).
11. M. Gilson, G. Zamora-López, V. Pallarés, *et al.*, "Model-based whole-brain effective connectivity to study distributed cognition in health and disease," Netw. Neurosci. **4**, 338–373 (2020).
12. M. Gilson, G. Deco, K. J. Friston, *et al.*, "Effective connectivity inferred from fmri transition dynamics during movie viewing points to a balanced reconfiguration of cortical interactions," NeuroImage **180**, 534–546 (2018). Brain Connectivity Dynamics.
13. G. Deco, C. Lynn, Y. S. Perl, and M. L. Kringelbach, "Violations of the fluctuation-dissipation theorem reveal distinct non-equilibrium dynamics of brain states," (2023).
14. M. L. Kringelbach, Y. S. Perl, E. Tagliazucchi, and G. Deco, "Toward naturalistic neuroscience: Mechanisms underlying the flattening of brain hierarchy in movie-watching compared to rest and task," Sci. Adv. **9**, eade6049 (2023).
15. L. F. Cugliandolo, J. Kurchan, and G. Parisi, "Off equilibrium dynamics and aging in unfrustrated systems," J. de Physique **4**, 1641–1656 (1994).
